# A Brownian ratchet model explains the biased sidestepping movement of single-headed kinesin-3 KIF1A

**DOI:** 10.1101/536250

**Authors:** A. Mitra, M. Suñé, S. Diez, J. M. Sancho, D. Oriola, J. Casademunt

## Abstract

The kinesin-3 motor KIF1A is involved in long-ranged axonal transport in neurons. In order to ensure vesicular delivery, motors need to navigate the microtubule lattice and overcome possible roadblocks along the way. The single-headed form of KIF1A is a highly diffusive motor that has been shown to be a prototype of Brownian motor by virtue of a weakly-bound diffusive state to the microtubule. Recently, groups of single-headed KIF1A motors were found to be able to sidestep along the microtubule lattice, creating left-handed helical membrane tubes when pulling on giant unilamellar vesicles *in vitro*. A possible hypothesis is that the diffusive state enables the motor to explore the microtubule lattice and switch protofilaments, leading to a left-handed helical motion. Here we study microtubule rotation driven by single-headed KIF1A motors using fluorescene-interference contrast (FLIC) microscopy. We find an average rotational pitch of ≃ 1.4 *μ*m which is remarkably robust to changes in the gliding velocity, ATP concentration and motor density. Our experimental results are compared to stochastic simulations of Brownian motors moving on a two-dimensional continuum ratchet potential, which quantitatively agree with the FLIC experiments. We find that single-headed KIF1A sidestepping can be explained as a consequence of the intrinsic handedness and polarity of the microtubule lattice in combination with the diffusive mechanochemical cycle of the motor.

## INTRODUCTION

Molecular motors from the kinesin and dynein family move along microtubules transducing the chemical energy of ATP hydrolysis into mechanical work (1). They perform a variety of mechanical functions in cells, which involve intracellular transport, flagellar beating or cytoplasmic streaming (1, 2). Microtubule filaments are hollow cylindrical structures consisting of several protofilaments providing a parallel array of tracks for the motors to move (3). Interestingly, motors do not always follow single protofilament tracks, but in some cases they are capable of switching protofilaments in a consistent manner (4–15). Such biased off-axis motion leads to helical trajectories of the motors with rotational pitches ranging from ∼ 0.3 to 3 *μ*m. This phenomenon seems to be general, and it has been reported for a great variety of motors such as monomeric kinesin-1 (4), kinesin-2 (5), kinesin-5 (6), kinesin-8 (7–9), kinesin-14 (10, 11), axonemal dynein (12–14) or cytoplasmic dynein (8, 15). Despite the numerous studies, the role and underlying mechanisms of such helical motions are still unclear. In the context of intracellular transport, it has been hypothesized that sidestepping might be a useful strategy to navigate the microtubule lattice and circumvent possible roadblocks (7, 16–20). What are the mechanisms enabling molecular motors to sidestep along the microtubule lattice in a biased manner and to generate torques? Diffusive search has been conjectured to be an important ingredient for such biased motion (4, 5), promoting lane-changing events (21–23), in contrast to the linear motion of motors that follow a single protofilament (e.g. dimeric kinesin-1 (24, 25)). Recently, the single-headed form of KIF1A (kinesin-3) was added to the list of motors that can generate helical movements (20). It was found that single-headed KIF1A motors were able to pull on membrane tubes and wind them around single microtubules *in vitro* (20). Several studies support the idea that the single-headed form of KIF1A acts as a prototype of Brownian motor by virtue of a weakly-bound diffusive state on the microtubule lattice (26, 27). This diffusive state makes the motor inefficient; however, it can lead to cooperative force generation (28, 29) and might be crucial for sidestepping on the microtubule lattice (20). However, the last hypothesis has not been carefully studied yet.

Here, we characterize the sidestepping motion of single-headed KIF1A by studying the longitudinal rotations of microtubules in gliding motility assays using fluorescence-interference contrast (FLIC) microscopy (8, 24). The average rotational pitch of microtubules is found to be ≃ 1.4 *μ*m and surprisingly, it is highly robust to changes in the gliding velocity, ATP concentration, microtubule length and motor density. This is in contrast to other recently studied kinesins such as Kip3/kinesin-8 (9) or Ncd/kinesin-14 (11), in which the rotational pitch is ATP dependent. In order to understand this phenomenon we performed numerical simulations of Brownian motors exploring the microtubule lattice (26–29). Such a simple model successfully reproduces the experimental observations in the FLIC experiments in terms of speed, rotational pitch and frequency of the gliding microtubules. We propose that the microtubule lattice geometry together with the microtubule-motor interaction, determines the rotational pitch.

## METHODS

### Protein expression and purification

Tubulin was purified from porcine brain (Vorwerk Podemus, Dresden, Germany) using established protocols as described previously (30). Single-headed KIF1A motor proteins (aminoacids 1-382) with both, a His-tag and a Cys residue at the N-terminus (27), were expressed in *Escherichia coli*, purified using a Ni-NTA column and labeled with biotin during elution, as described in Ref. (20).

### Polymerization of speckled microtubules

Polymerization of speckled microtubules Guanylyl-(*α, β*)-methylene-diphosphonate (GMP-CPP)-grown, taxol-stabilized rhodamine-speckled microtubules (’speckled microtubules’) were grown as described in Ref. (8).

### KIF1A gliding assays on silicon wafers

The gliding assays were performed in microfluidic flow chambers constructed by sealing 22 mm × 22 mm glass coverslips (Menzel, Braunschweig, Germany; #1.5) and 10 mm × 10 mm silicon wafers having a 30 nm thermally grown oxide layer (GESIM, Grosserkmannsdorf, Germany) with spacers of NESCO film (Azwell Inc., Osaka, Japan). Both, glass and wafer surfaces, were coated with dichlorodimethylsilane to render the surface hydrophobic (31). The approximate dimension of each flow chamber was 10 mm × 1.5 mm × 100 *μ*m. Flow chambers were flushed with solutions (15 *μ*l each) in the following sequence: (1) fab-fragment solution consisting of 100 *μ*g/ml F(ab’)2 fragments (anti-mouse IgG [Fc specific] antibody developed in goat; Sigma) in PBS that bind unspecifically to the hydrophobic surface (incubation time 5 min), (2) Pluronic F127 (Sigma, 1% in PBS) in order to block the surface against unspecific protein adsorption (incubation time > 60 min), (3) antibody solution consisting of 100 *μ*g/ml biotin tubulin antibodies (Thermo Fisher Scientific) in PBS that bind specifically to the F(ab’)2 fragments (incubation time 5 min), (4) four times BRB40 buffer (40 mM Pipes [Sigma], pH 6.84, with KOH [VWR], 1mM EGTA [Sigma], 4mM MgCl_2_ [VWR]) to remove excess F-127 in solution and exchange buffers, (5) motor solution consisting of 2 *μ*M biotinylated KIF1A in motor dilution buffer (1mM ATP [Roche], 0.1% Tween 20 [Sigma], 0.2 mg/ml casein [Sigma] and 0.2 mg/ml DTT [Sigma] in BRB40) in order to bind the KIF1A proteins specifically to the biotin antibodies (incubation time 5 min), (6) speckled-microtubule solution consisting of speckled microtubules in motor dilution buffer (incubation time 5 min), and (7) imaging solution containing the oxygen scavenger system (40 mM glucose [Sigma], 110 *μ*g/ml glucose oxidase [SERVA], 22 *μ*g/ml catalase [Sigma]) in motor dilution buffer to remove excess microtubules. The gliding assay scheme is illustrated in Fig. 1A.

**Figure 1:**
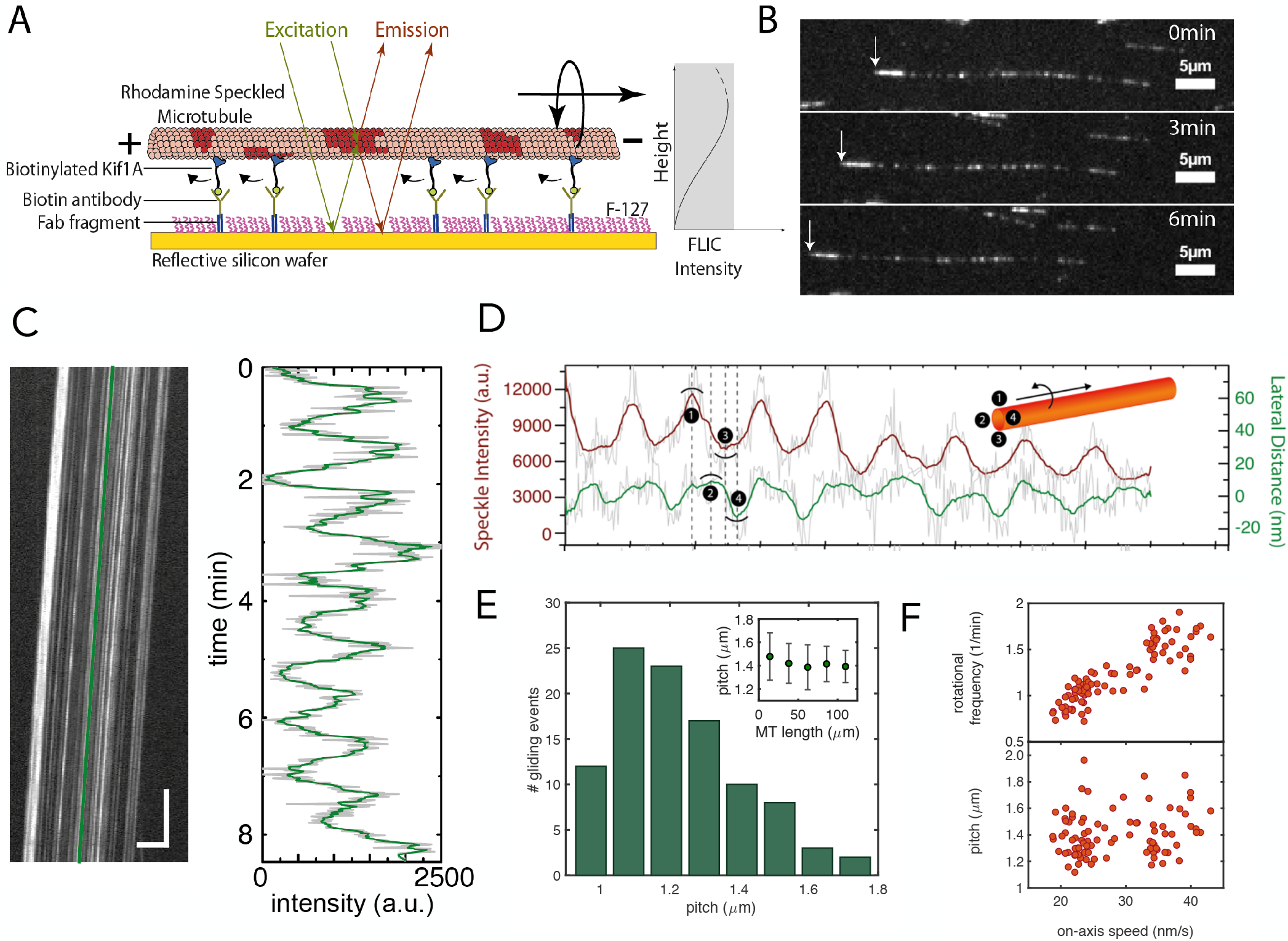
Rotational motion of speckled microtubules gliding on single-headed KIF1A. (A) Schematic representation of a speckled microtubule gliding on a reflective silicon substrate coated with biotinylated KIF1A motors via biotin antibodies. (B) Fluorescent image series of an example rhodamine speckled microtubule (minus end marked by white arrow) gliding with a velocity of 22 nm/s (see Supplementary Movie 1). (C) Corresponding kymograph (space-time intensity plot) of the speckled microtubule in (B) (horizontal scale bar: 5 *μ*m; vertical scale bar: 1 min) and the FLIC intensity profile over time for one of the speckles indicated by the green line in the kymograph. The rotational pitch for this microtubule was 1.1 *μ*m. (D) Direction of rotation of the microtubules gliding on KIF1A: An individual speckle from a gliding speckled microtubule was tracked using FIESTA to obtain the lateral deviation of the speckle along with the variation in FLIC intensity over time. Raw data is indicated in light grey and the smoothened data (rolling frame averaged over 20 frames) is indicated in green (lateral distance; positive values refer to left) and brown (FLIC intensity). Inset: Illustration of the counterclockwise rotation of a microtubule in the direction of motion. (E) Histogram of rotational pitches showing a median pitch of 1.4 ± 0.2 *μ*m (median ± SD, *n* = 100 gliding events). Inset: Variation of rotational pitch with respect to microtubule length by binning the data (mean ± SD). (F) The rotational frequency (top) and rotational pitch (bottom) plotted with respect to the on-axis microtubule gliding velocity.

### Image acquisition

Optical imaging was performed using an inverted fluorescence microscope (Zeiss Axiovert 200M, Carl Zeiss, Göttingen, Germany) with a 63× water immersion 1.2 NA objective (Zeiss) in combination with an Andor Ixon DV 897 (Andor Technology, Belfast, UK) EMCCD camera controlled by Metamorph (Molecular Devices Corporation, Sunnyvale, CA, USA) providing a pixel size of 0.256 *μ*m. A Lumen 200 metal arc lamp (Prior Scientific Instruments Ltd., Cambridge, UK) was used for epifluorescence excitation. Speckled microtubules gliding on the surface of the silicon-wafer were imaged ‘through the solution’ (i.e. on the far side of the flow channels) using a TRITC filterset (Ex 534/30×, DC BC R561, EM BL593/40, all Chroma Technology Corp., Rockingham, VT) with an exposure time of 400 ms per frame. Images were recorded in time-lapse mode with a frame rate of 1 fps.

### Image analysis

Kymographs (space-time intensity plots) for gliding speckled microtubules were generated in ImageJ (32). The kymographs were then analysed with MATLAB (Mathworks, Natick, MA) using the speckle analysis method described in (8) to obtain the rotational pitch and velocity corresponding to each gliding speckled microtubule. To obtain information regarding the direction of rotation, example speckles were tracked using FIESTA (33) and the tracks were averaged (rolling frame window of 2 *μ*m) to get the estimated microtubule centerline. The perpendicular distance of a speckle track from the microtubule centerline at each point provided the lateral deviation of the speckle. In combination with the FLIC intensity variation, this information provided the direction of rotation (see Fig. 1D and Supplementary Figure 2).

### Simulation details

KIF1A trajectories were simulated by solving Eq. 3 from the Supplementary Material using a second–order Euler method (Heun’s method), including a two-dimensional potential landscape given by Eq. 2 from the Supplementary Material. The numerical simulation of the stochastic part was carried out through a Gaussian random variate with null mean and variance equal to 1 according the procedure developed in (34). To generate random numbers, we used the Mersenne Twister generator and the alternative *Marsaglia-Tsang ziggurat* and *Kinderman-Monahan-Leva* ratio methods (35). The discretization time step Δ*t* was chosen as the smallest time scale of the system. *In vitro* experiments and previous studies (20, 26–28, 36) provided values for several parameters used in the simulation: *D* = 20 nm^2^ ms^−1^, *τ* = 4 ms, *τ^*^* = 4–42 ms, *θ* = 0.45*π*, *V*_0_ = 20 *k_B_T*, and *U^*^* = 2*V*_0_/100.

## RESULTS

### Experimental results

To investigate the microtubule rotation driven by single-headed KIF1A motors we performed gliding motility assays by (1) coating a reflective silicon surface with biotinylated KIF1A motors and (2) flushing in rhodamine speckled microtubules (illustrated in Fig. 1A). Biotinylated motors were linked to the surface via biotin antibodies, which in turn were bound to F(ab’)2 (or Fab) fragments coating the surface. Microtubules glided robustly with constant velocities in the range of 20 – 40 nm/s (see Supplementary Movie 1 and Fig. 1B). The microtubule gliding velocity did not vary with the concentration of Fab fragments, indicating no dependence with the motor density (Supplementary Figure 1). This result is in agreement with theoretical studies predicting an early saturation of the speed of KIF1A clusters with motor number (29). The reflective surface of the gliding assay enabled us to use FLIC microscopy to obtain rotational information in combination with forward motion for the gliding microtubules (7, 8, 24). As seen in Fig. 1C, the recorded intensities of the speckles fluctuated periodically due to changes in speckle height with respect to the reflective surface. The periodicity of the intensity fluctuation provided us with the rotational pitch of the gliding microtubules. The mean rotational pitch was 1.4 ± 0.2 *μ*m (mean ± SD, *n* = 100 gliding events, see Fig. 1E) much shorter than the supertwist of GMP-CPP grown taxol stabilized microtubules (∼ 8 *μ*m) (8, 24). To obtain the direction of rotation, several individual speckles from different microtubules were tracked using the tracking software FIESTA (33). In combination with the height information (obtained by FLIC), the sideways information revealed that the microtubules rotated counterclockwise in the direction of gliding motion (see Fig. 1D and additional examples in Supplementary Figure 2). The rotational pitch did not vary significantly as a function of microtubule length (Fig. 1E, inset). Furthermore, there was no significant dependence of the rotational pitch on the gliding velocity (Fig. 1F, bottom). This implies that the rotational frequency, i.e., the number of motor sidesteps on the microtubule lattice per unit time, increases with increasing gliding velocity (Fig. 1F, top). In order to investigate more carefully the dependence of the rotational pitch on the motor speed, we varied the gliding velocity by changing the ATP concentration. Interestingly, we observed that a decrease in the ATP concentration reduced the gliding velocity but not the rotational pitch (Supplementary Figure 3). Hence, we conclude that the rotational pitch is remarkably robust to changes on the gliding velocity, ATP concentration, microtubule length and motor density.

### Simulation results

In order to understand how KIF1A motors select the rotational pitch, we performed stochastic simulations of the dynamics of single KIF1A motors. Based on experimental observations and theoretical studies (20, 26–29), we used a two-state Brownian ratchet model to simulate the dynamics of the motor on the microtubule lattice (see Fig. 2A-D and Supplementary Material).

**Figure 2:**
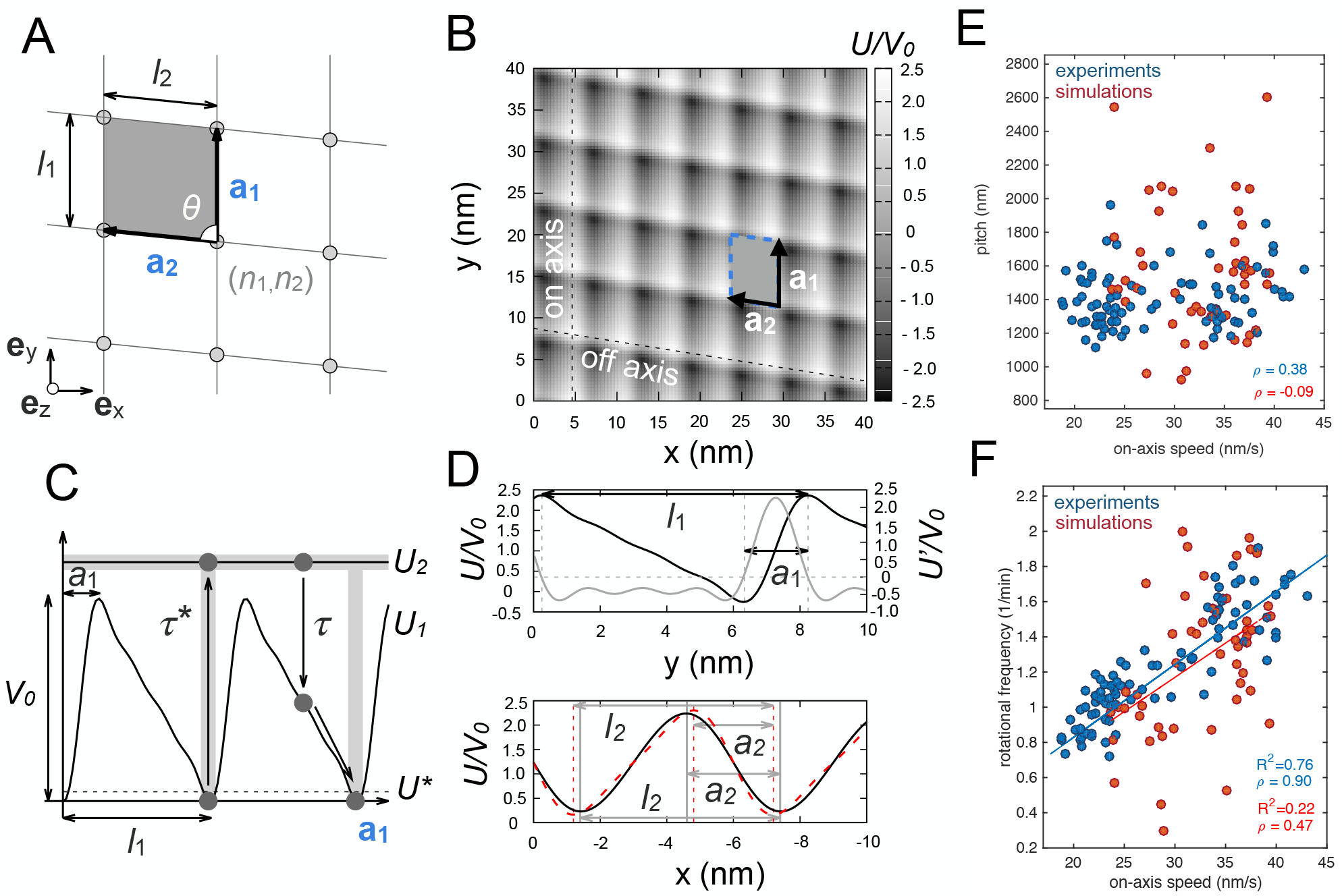
A) Oblique Bravais lattice as a description of the microtubule lattice with primitive vectors **a**_1_ and **a**_2_ of size *l*_1_ and *l*_2_ respectively, forming an angle *θ*. The gray parallelogram corresponds to the primitive cell of the lattice and the gray circles to the nodes of the lattice. B) Two–dimensional microtubule-motor potential with *N*_1_ = 4, *N*_2_ = 2 and coefficients *μ*_11_ = 1, *μ*_12_ = 0.9, *μ*_13_ = 0.65, *μ*_14_ = 0.35, *μ*_21_ = 1, and *μ*_22_ = 0.085 (see Supplementary Information). Dashed lines–directions along which 1*d* sections of the potential are plotted in (D). C) Sawtooth linear potential along **a**_1_ (*N*_1_ = 4, *μ*_11_ = 1, *μ*_12_ = 0.9, *μ*_13_ = 0.65, *μ*_14_ = 0.35) for the motor–track interaction. The gray regions depict the zones where excitations from *U*_1_ to *U*_2_ are allowed with exponentially distributed hydrolysis dwell times with mean *τ^*^*. Transitions from *U*_2_ to *U*_1_ are delocalized and occur with exponentially distributed decay times with mean *τ*. Dashed line–excitation time starts when the particle’s potential energy is lower than *U^*^*. D) Top: Potential section along **a**_1_. Gray–partial derivative of the potential, its roots (intersections of the gray dashed lines) label the maxima and minima of the potential. *l*_1_ = 7.9 nm, *a*_1_ = 1.9 nm (*a*_1_*/l*_1_ = 0.24). Bottom: Potential section along **a**_2_; *l*_2_ = 6.0 nm, *a*_2_ = 2.8 nm (*a*_2_*/l*_2_ = 0.47). Red—potential section along **a**_2_ when *μ*_22_ = 0.4 (the rest being the same); *l*_2_ = 6.0 nm, *a*_2_ = 2.4 nm (*a*_2_*/l*_2_ = 0.4). E) Rotational pitch and F) the corresponding rotational frequency for the simulation results (red circles) and the experimental data from Fig. 1F (blue circles). The simulations are obtained for an ensemble of 10^3^ independent trajectories at zero load using a landscape generated according to Eq. 2 from the Supplementary Material with asymmetry *a*_1_*/l*_1_ = 0.24, *a*_2_*/l*_2_ = 0.47 and varying the force applied in the direction of motion for *τ^*^* = 46 ms (see Supplementary Material). The correlation of the rotational pitch and the rotational frequency with the on-axis speed is provided by the Pearson correlation coefficient *ρ*. The lines correspond to fitted linear regressions with their respective *R*^2^ values.

In order to test the robustness of the pitch to changes on the gliding velocities, we studied the pitch and the rotational frequency for different gliding velocities to compare the simulations with the experimental results in Fig. 1F. There exist two main ways to change the gliding velocity in the simulations: (i) varying the ATP hydrolysis dwell time at zero load or (ii) applying a certain force along the direction of motion. In the first case, we obtained that the pitch did not vary significantly as a function of the speed of the motors, while the rotational frequency increased approximately linearly (see Supplementary Figure 4C,D) in qualitative agreement with the experiments by changing the ATP concentration (Supplementary Figure 3).

In the second case, we changed the load along the direction of motion and we kept the same value for the dwell time. This scenario mimics possible friction forces experienced by the motors during the gliding assay that might cause a dispersion on the velocities. The results obtained on the rotational pitch and frequency followed similar trends as for varying dwell times (Supplementary Figure 3 and 4C,D), and they quantitatively agreed with the experimental results (see Figure 2 E,F). Altogether, we conclude that a Brownian ratchet description of single-headed KIF1A successfully explains the essential features observed experimentally.

## DISCUSSION

In this work we have shown that the rotation of microtubules due to the action of single-headed KIF1A motors can be explained by a simple Brownain ratchet model on the microtubule lattice. Interestingly, the rotational pitch is independent on the gliding velocity which implies that the frequency of rotation increases linearly with the speed. These results are fundamentally different from recent studies on Kip3/Kinesin-8 or ncd/kinesin-14, where the rotational pitch depends on the gliding velocity (9, 11). In the latter cases, an increase on the dwell time to hydrolyze an ATP molecule favours sidestepping respect to forward stepping, whereas in our case the dwell time affects both on-axis and off-axis movements, leaving the pitch unaffected. The rotational pitch is found to be very similar to the one observed in tube pulling experiments (20), where KIF1A motors formed helical tubes around microtubules. This implies that both weak (membrane-bound) and strong (glass-bound) anchoring lead to similar results for the selection of the pitch. Our numerical simulations can reproduce such pitch values considering almost no lateral asymmetry (*a*_2_ ≃ 0.47). This asymmetry parameter is likely to depend on the microtubule-motor interaction and thus be characteristic of each kinesin type. Slightly larger asymmetries (*a*_2_ ≃ 0.4) lead to shorter pitch values (≃ 300 nm) which could account for the observed pitch of other diffusive motors such as single-headed kinesin-1 (4, 6). In this work, we cannot rule out the possibility of collective effects affecting the observed experimental pitch. One possibility is that the measured pitch reflects the averaged pitch of a single motor. In this case, collective effects are not important and groups of motors would only reduce the diffusivity of the rotational movement, leaving the pitch unaffected. A second possibility is that the single motor pitch is small but collective effects play an important role and select a larger pitch, as proposed in Ref. (20). Our results are consistent with the first option, although they do not discard the second option. In order to discern between these two possibilities, single-molecule tracking of the helical movement of single-headed KIF1A should be performed, similarly as in Ref. (9). Finally, our model can potentially explain the pitch of other weakly processive motors such as single-headed kinesin-1 or kinesin-5 by changing the lateral asymmetry. Further work needs to be undertaken to confirm if the present mechanism also applies to other kinesin motors.

## Authors’ contributions

A.M. performed experimental research and M.S. and D.O. performed theoretical research. D.O. and J.C. conceived the project and J.-M. S., S.D., D.O. and J.C. directed the project. A.M, M.S. and D.O wrote the manuscript. All authors approved the final version of the manuscript.

## Acknowledgements

We are grateful to N. Hirokawa (University of Tokyo) for kindly providing us the KIF1A construct A382 (27). We thank M. Gironella for her contribution on theoretical modelling at the early stage of the project.

## References

1. Howard, J. 2001. Mechanics of motor proteins and the cytoskeleton. Sinauer Associates Sunderland, MA.

2. Alberts B., A. Johnson, J. Lewis, M. Raff, K. Roberts and P. Walter. 1994. Molecular Biology of the Cell. Garland, New York.

3. Amos, L. A, and D. Schlieper. Microtubules and MAPs. 2005. Advances in protein chemistry. 71:257–298.

4. Yajima, J., R. A. Cross. A torque component in the kinesin-1 power stroke. 2005. Nat. Chem. Biol. 1:338–341.

5. Brunnbauer, M., R. Dombi, T.-H. Ho, M. Schliwa, M. Rief, Z. Ökten. Torque generation of kinesin motors is governed by the stability of the neck domain. 2012. Mol. Cell. 46, 147–158.

6. Yajima, J., K. Mizutani and T. Nishizaka. A torque component present in mitotic kinesin Eg5 revealed by three-dimensional tracking. 2008. Nat. Struct. Mol. Biol. 15:1119 – 1121.

7. Bormuth, V., B. Nitzsche, F. Ruhnow, A. Mitra, M. Storch, B. Rammner, J. Howard and S. Diez. The highly processive kinesin-8, Kip3, switches microtubule protofilaments with a bias toward the left. Biophys. J. 2012. 103:L4–L6.

8. Mitra, A., F. Ruhnow, B. Nitzsche and S. Diez. Impact-free measurement of microtubule rotations on kinesin and cytoplasmic-dynein coated surfaces. 2015. PLoS One. 10:e0136920. (doi:10.1371/journal.pone.0136920).

9. Mitra, A., F. Runhow, S. Girardo, S. Diez. Directionally-biased sidestepping of Kip3/kinesin-8 is regulated by ATP waiting time and motor-microtubule interaction strength. 2018. Proc. Natl. Acad. Sci. U.S.A. 115:E7950–E7959.

10. Walker, R. A., E. D. Salmon and S.A. Endow. The Drosophila claret segregation protein is a minus-end directed motor molecule. Nature. 1990. 347:780–782.

11. Nitzsche, B., E. Dudek, A. A. Hajdo, A. Kasprzak, A. Vilfan and S. Diez. The working stroke of the kinesin-14, ncd, comprises two substeps of different direction. 2016. Proc. Natl. Acad. Sci. U.S.A. 113: E6582–E6589.

12. Yamaguchi, S., K. Saito, M. Sutoh, T. Nishizaka, Y. Toyoshima and J. Yajima. Torque Generation by Axonemal Outer-Arm Dynein. 2015. Biophys. J.,108: 872–879.

13. Vale, R. and Y. Toyoshima. Rotation and translocation of microtubules in vitro induced by dyneins from Tetrahymena cilia. 1988. Cell. 52:459–469.

14. Mimori, Y. and T. Miki-Noumura. Extrusion of rotating microtubules on the dynein-track from a microtubule-dynein gamma-complex. 1995. Cell Motil. Cytoskel. 30:17–25.

15. Can, S., M. A. Dewitt and A. Yildiz. Bidirectional helical motility of cytoplasmic dynein around microtubules. 2014. Elife. e03205.

16. Bugiel, M., E. Böhl, E. Schäffer. The Kinesin-8 Kip3 Switches Protofilaments in a Sideward Random Walk Asymmetrically Biased by Force. 2015. Biophy J.108:2019–2027.

17. Hoeprich, G.J., A. R. Thompson, D. P. McVicker, W. O. Hancock and C. L. Berger. Kinesin’s neck-linker determines its ability to navigate obstacles on the microtubule surface. 2014. Biophy J. 106:1691–1700.

18. Hoeprich, G.J., K. J. Mickolajczyk, S.R. Nelson, W. O. Hancock and C. L. Berger. The axonal transport motor kinesin-2 navigates microtubule obstacles via protofilament switching. 2017. Traffic. 18:304–314.

19. Schneider, R., T. Korten, W. J. Walter and S. Diez. Kinesin-1 Motors Can Circumvent Permanent Roadblocks by Side-Shifting to Neighboring Protofilaments. 2015. Biophys. J. 108:2249–2257.

20. Oriola, D., S. Roth, M. Dogterom and J. Casademunt. Formation of helical membrane tubes around microtubules by single-headed kinesin KIF1A. 2015. Nat. Commun. 6:8025 (doi: 10.1038/ncomms9025).

21. Chowdhury, D., A. Garai and J.-S. Wang. Traffic of single-headed motor proteins KIF1A: Effects of lane changing. 2008. Phys. Rev. E. 77:050902(R).

22. Curatolo, A. I., M. R. Evans, Y. Kafri and J. Tailleur. Multilane driven diffusive systems. 2016. J. Phys. A: Math. Theor. 49: 095601.

23. Verma, A. K. and A. K. Gupta. Effect of Binding Constant on Phase Diagram for Three-Lane Exclusion Process. 2016. Adv. Comp. and Comm. Tech. 289–296

24. Nitzsche, B., F. Ruhnow and S. Diez. Quantum-dot assisted characterization of microtubule rotations during cargo transport. 2008. Nat. Nanotechnol. 2008.1: 1–9.

25. Ray, S., E. Meyhöfer, R. A. Milligan and J. Howard. Kinesin follows the microtubules protofilament axis. 1993. J Cell Biol. 121:1083–1093.

26. Okada, Y. and N. Hirokawa. A processive single-headed motor: kinesin superfamily protein KIF1A. 1999. Science. 283:1152–1157.

27. Okada, Y., H. Higuchi, N. Hirokawa. Processivity of the single-headed kinesin KIF1A through biased binding to tubulin. 2003. Nature. 424:574–577.

28. Oriola, D. and J. Casademunt. Cooperative force generation of KIF1A Brownian motors. 2013. Phys. Rev. Lett. 111:048103.

29. Oriola, D. and J. Casademunt. Cooperative action of KIF1A Brownian motors with finite dwell time. 2014. Phys. Rev. E. 89:032722.

30. Castoldi, M. and A. V. Popov. Purification of brain tubulin through two cycles of polymerization-depolymerization in a high-molarity buffer. Protein Expr. Purif. 2003. 32: 83–8. (doi:10.1016/S1046-5928(03)00218-3).

31. Korten, T., B. Nitzsche, C. Gell, F. Ruhnow, C. Leduc and S. Diez. Fluorescence imaging of single Kinesin motors on immobilized microtubules. 2011. Methods Mol. Biol. 2011. 783: 121–37. (doi:10.1007/978-1-61779-282-37)

32. Schneider, C.A., W. S. Rasband, K. W. Eliceiri. NIH Image to ImageJ: 25 years of image analysis. Nat. Methods. 2012. 9:671–675.

33. Ruhnow, F., D. Zwicker and S. Diez. Tracking single particles and elongated filaments with nanometer precision. Biophys. J. 2011.100:2820–8. (doi:10.1016/j.bpj.2011.04.023)

34. Garcia-Ojalvo, J. and J.-M. Sancho. Noise in spatially extended systems. 1999. Springer, New York.

35. Galassi, M., J. Davies, J. Theiler, B. Gough, G. Jungman, P. Alken, M. Booth and F. Rossi. GNU Scientic Library Reference Manual. 2009. Network Theory Ltd.

36. Nishinari, K., Y. Okada, A. Schadschneider and D. Chowdhury. Intracellular transport of single-headed molecular motors KIF1A. 2005. Phys. Rev. Lett. 95:118101.

