## Supplementary Material for "A Brownian ratchet model explains the biased sidestepping movement of single-headed kinesin-3 KIF1A"

### Two-dimensional Brownian motor model for single-headed KIF1A

We define the position of a motor in Cartesian coordinates as $\mathbf{r} = x\,\mathbf{e}_x + y\,\mathbf{e}_y$ , where $\mathbf{e}_y$ corresponds to the microtubule axis direction (see Fig. 2A). The microtubule lattice is treated as an oblique Bravais lattice with primitive vectors $\mathbf{a}_i$, $i = 1, 2$, forming an angle $\theta > 0$ (see Figs. 2A,B). Hereinafter, latin indices will be employed to label the primitive directions, and greek indices to label the Cartesian coordinates. The nodes of the lattice are defined as $\mathbf{R}(n_1, n_2) = n_1\mathbf{a}_1 + n_2\mathbf{a}_2$ where $n_1, n_2$ denote integer numbers and correspond to the available binding sites for the motor. Hence, the set $(n_1, n_2)$ indicates a given primitive cell. The two primitive vectors are $\mathbf{a}_1 = l_1\,\mathbf{e}_y$ and $\mathbf{a}_2 = l_2(-\sin\theta\,\mathbf{e}_x + \cos\theta\,\mathbf{e}_y)$, where $l_i$, $i = 1, 2$ are the periodicities for each primitive direction (see Figs. 2A,B). The reciprocal basis $\mathbf{q}_i$ reads:

$$\mathbf{q}_1 = 2\pi\,\frac{\mathbf{a}_2 \times \mathbf{e}_y}{|\mathbf{a}_1 \times \mathbf{a}_2|} = \frac{2\pi}{l_1}\,(\cot\theta\,\mathbf{e}_x + \mathbf{e}_y); \quad \mathbf{q}_2 = 2\pi\,\frac{\mathbf{e}_y \times \mathbf{a}_1}{|\mathbf{a}_1 \times \mathbf{a}_2|} = -\frac{2\pi}{l_2}\,\csc\theta\,\mathbf{e}_x\,. \tag{1}$$

We consider the motor can be in two possible states $k = 1, 2$. State $k = 1$ corresponds to a 'strongly–bound' state, in which the motor experiences a potential landscape $U_1(x, y)$, describing the microtubule-motor interaction (see Fig. 2C). In this state the motor is not bound to an ATP molecule. After capturing an ATP molecule, hydrolysis takes place and the motor excites to state $k = 2$ usually known as the 'weakly–bound' (or ADP-P$_i$) state, in which the motor can freely diffuse on the lattice (see Fig. 2C). In this state the motor experiences a flat potential. After the release of the ADP and P$_i$ molecules, the motor decays from state $k = 2$ to the state $k = 1$. We consider this process occurs with exponentially distributed decay times with mean $\tau$. Decay processes are de-localized in space, whereas excitations are localized in the lattice nodes ($U_1$ potential minima). A transition to the weakly–bound state $k = 2$ is only allowed if the potential energy in state 1 is below a certain threshold value $U^\star$, much smaller than $U_1$. Again, an exponential distribution of exciting times is assumed, with mean time $\tau^*$. The potential $U_1(x, y)$ must be asymmetric in the $y-$component, reflecting microtubule polarity. A common choice is a sawtooth landscape potential. The reciprocal basis $\{\mathbf{q}_1, \mathbf{q}_2\}$ is employed to build a periodic two-dimensional oblique sawtooth landscape, with periodicities $l_1$, $l_2$ along the primitive directions $\mathbf{a}_1$, $\mathbf{a}_2$, respectively. A simple choice is to define $U_1 = V_1 + V_2$, with $V_i$, $i = 1, 2$ being one–dimensional sawtooth functions in each primitive direction. The one–dimensional sawtooth functions may be expressed as Fourier series,

$$V_i(x, y) = V_0 \sum_{j=1}^{N_i} \frac{\mu_{ij}}{j}\,\sin(j\mathbf{q}_i \cdot \mathbf{r})\ ;\ i = 1, 2 \tag{2}$$

A simple choice of the coefficients $\mu_{ij}$ leads to a continuous smooth landscape (see Figs. 2B,C,D). The two-dimensional ratchet may have two distinct asymmetries $a_i$ along the directions $\mathbf{a}_i$, $i = 1, 2$ (see Figs. 2C,D). As previously mentioned, the polarity of a microtubule generates an asymetry $a_1$ in the $\mathbf{a}_1$ direction which has been estimated to be $a_1/l_1 \simeq 0.2$ (1). In general, a second asymmetry $a_2/l_2 \neq 1/2$ can exist along the $\mathbf{a}_2$ direction. However, even in the symmetric case ($a_2/l_2 = 1/2$), the fact that $\theta \neq \pi/2$ and $a_1/l_1 \neq 1/2$ is sufficient to generate a helical movement of the motor (data not shown). From Eq. 2, wee see that the asymmetry is not a parameter itself in our model, but it is determined by the choice of the coefficients $\mu_{ij}$. Two harmonics may be enough to generate an asymmetric landscape; however, the choice of $a_i$ is limited because for $\mu_{i2} \geq 1$ the potential displays a non-desirable second local minimum for each period. This problem can be overcome by using higher order harmonics as shown in Figs. 2C,D. In order to match the experimental results, we choose the asymmetry parameters $a_1/l_1 = 0.24$ and $a_2/l_2 = 0.47$. The dynamics of the motor follows Langevin dynamics in the overdamped limit,

$$\eta\,\dot{\mathbf{r}} = -\delta_{k1}\,\nabla U_1 - \mathbf{F} + \boldsymbol{\xi}(t). \tag{3}$$

where $\eta$ is the friction coefficient fulfilling the Einstein's relation $D = k_B T/\eta$ ($D$ is the diffusion coefficient), the nabla operator reads $\nabla \equiv (\partial_x, \partial_y)$, $\mathbf{F}$ is a constant external force, and $\boldsymbol{\xi}$ is a random force vector with zero mean satisfying $\langle \xi_\alpha(t)\, \xi_\beta(t') \rangle = 2k_B T\, \eta\, \delta_{\alpha\beta}\, \delta(t-t')$, $\alpha, \beta = x, y$ where $\delta_{k1}$, $\delta_{\alpha\beta}$ are Kronecker deltas. Finally, we define a useful quantity to study the randomness of the motor motion on the lattice (2, 3),

$$R_\alpha = \frac{\langle \Delta r_\alpha^2(t) \rangle - \langle \Delta r_\alpha(t) \rangle^2}{d_\alpha \langle \Delta r_\alpha(t) \rangle}; \quad \alpha = x, y \tag{4}$$

where $\Delta r_\alpha(t) = r_\alpha(t) - r_\alpha(0)$ is evaluated at steady state, $d_\alpha$ is the motor step size —$d_x = l_2 \sin(\theta)$ along the $x$-axis, $d_y = l_1$ along the $y$-axis —, and the angle brackets denote an ensemble average. The parameter $R_\alpha$ quantifies the temporal irregularity of the dynamics of the motor along the $\alpha$-axis. A perfect clocklike motor would display no stepping irregularity, $R = 0$, whereas a motor undergoing exponentially distributed steps would exhibit $R = 1$ (3).

### Analysis of the rotational pitch and velocity of Brownian KIF1A motors under load

Single KIF1A trajectories are found to be very noisy and follow a diffusive behaviour at long time scales (Supplementary Figure 5). The motor movement is much more persistent in the longitudinal direction than in the transversal direction ($R_x \simeq 300$, $R_y \simeq 10$ see Supplementary Material). The rotational pitch of a trajectory $P$ and the frequency of rotation $\omega$ can be defined as $P = v_y\, t = v_y/\omega$ and $\omega = v_x/(2\pi R_{MT})$, where $R_{MT} = 12.5$ nm is the microtubule radius and $v_x, v_y$ are the speeds in the transversal and longitudinal directions, respectively. Considering parameters extracted from the literature (see Simulation details), at zero load we obtain a good agreement with experiments on the speed and stall force of a single KIF1A motor (1, 4) as well as on the rotational pitch (see Supplementary Figure 4A). Under on-axis loading ($\mathbf{F} = -F\, \mathbf{a}_1$), the velocity–force curve remains unaltered for changes in the lateral asymmetry $a_2$, whereas the pitch exhibits a strong dependence (see Supplementary Figure 4A). Finally, under off-axis loading ($\mathbf{F} = -F\, \mathbf{a}_2$) small changes in the lateral asymmetry have dramatic effects on the velocity-force relationship (Supplementary Figure 4B).

### References


1. Oriola, D. and J. Casademunt. Cooperative force generation of KIF1A Brownian motors. 2013. Phys. Rev. Lett. 111:048103.
2. Visscher, K., M. J. Schnitzer and S. M. Block. Single kinesin molecules studied with a molecular force clamp. 1999. Nature. 400:184–189.
3. Kolomeisky, A. B. and M. E. Fisher. Molecular Motors: A Theorist's Perspective. Ann. Rev. Phys. Chem. 2007. 58:675–695.
4. Okada, Y., H. Higuchi, N. Hirokawa. Processivity of the single-headed kinesin KIF1A through biased binding to tubulin. 2003. Nature. 424:574–577.

### SUPPLEMENTARY FIGURES

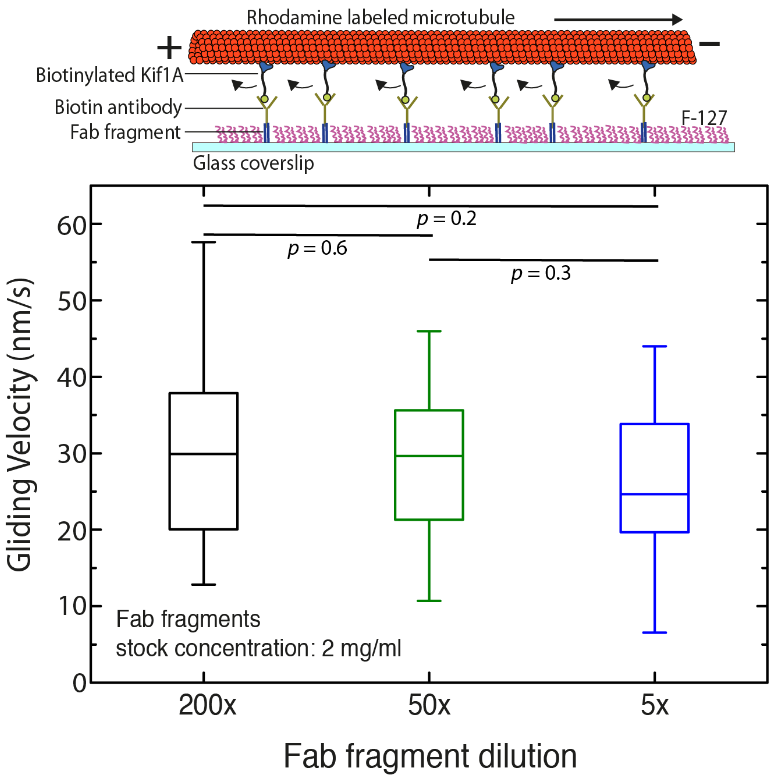


Supplementary Figure 1: Gliding velocity of microtubules as a function of the density of Fab fragments. The velocity of microtubules gliding on KIFA does not vary significantly with density of KIFA motors attached to the substrate. The figure shows box plots of microtubule gliding velocity for different Fab fragment dilutions (200x, 50x and 5x: stock concentration – 2mg/ml). The Fab fragment concentration determines the KIFA motor density on the surface.

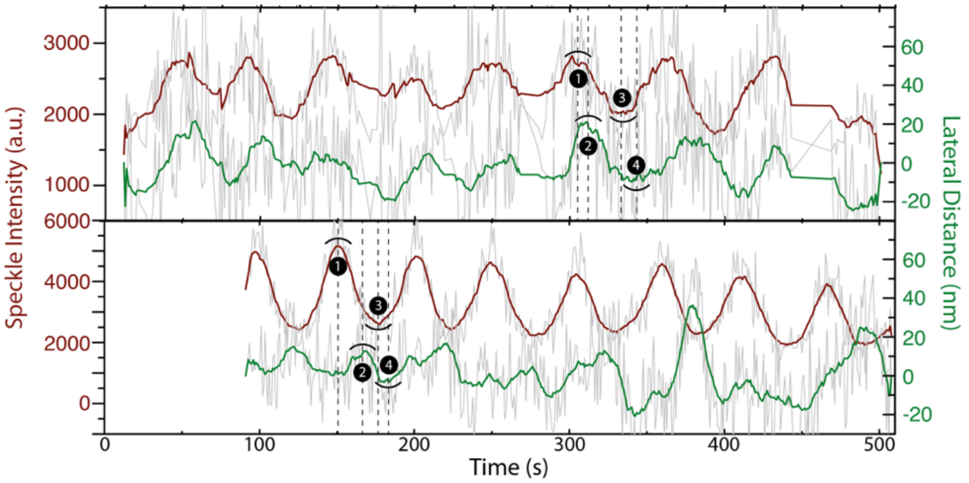


Supplementary Figure 2: Direction of rotation of the microtubules gliding on KIF1A. Two individual speckles from two gliding speckled microtubules were tracked using FIESTA to obtain the lateral deviation of the speckle along with the variation in FLIC intensity over time. The direction of rotation was determined to be counterclockwise (in the direction of motion) from the temporal sequence of the 3D position of the speckle. Raw data is indicated in light grey and the smoothened data (rolling frame averaged over 20 frames) is indicated in green (lateral distance; positive values refer to left) and brown (FLIC intensity). The numbers correspond to different positions on the microtubule surface (see Figure 1D, inset).

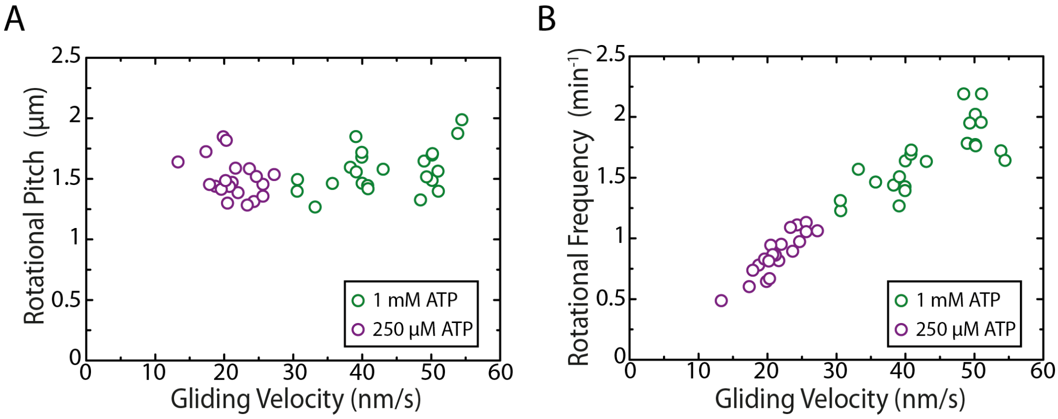


Supplementary Figure 3: Rotational pitch (A) and rotational frequency (B) plotted with respect to the gliding velocity for speckled microtubules gliding under low (250mM) and high (1mM) ATP conditions. The ATP concentration was switched in the same channel to keep all other parameters constant. A four-fold decrease in ATP concentration leads to an approximate two fold decrease of the gliding speed, with a subsequent reduction on the frequency of rotation but no effect on the rotational pitch.

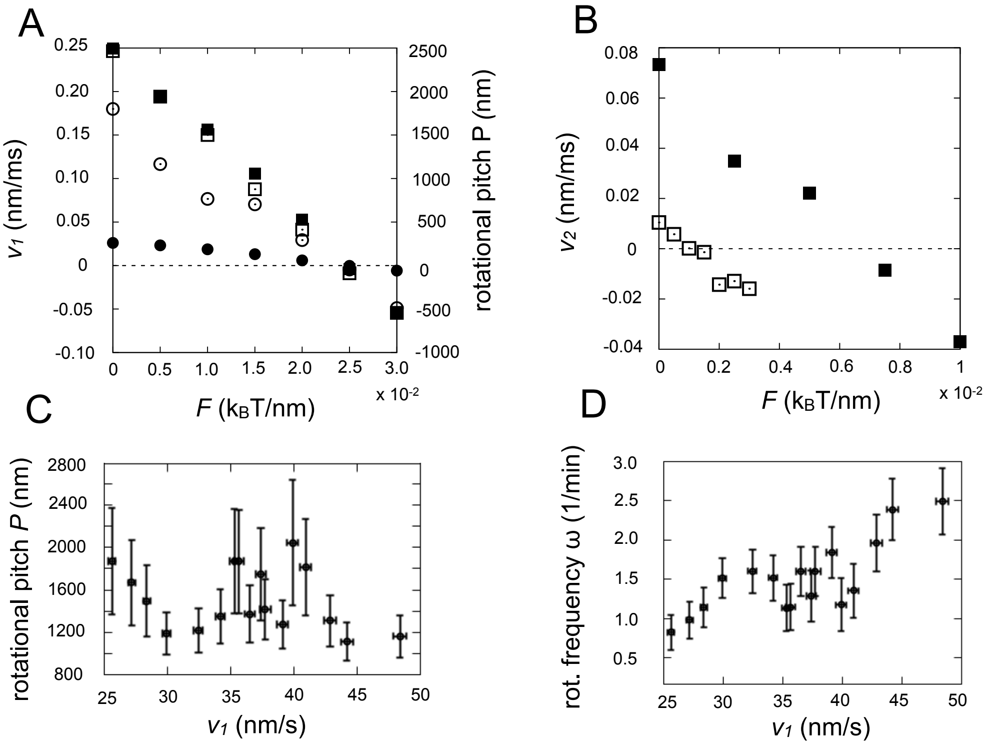


Supplementary Figure 4: A) Mean longitudinal velocity—squares—and rotational pitch—circles—for an ensemble of $10^3$ independent trajectories simulated in a landscape generated by Eq. (2) with asymmetries $a_1/l_1 = 0.24$, $a_2/l_2 = 0.47$—blank symbols—, and $a_1/l_1 = 0.24$, $a_2/l_2 = 0.40$—black filled symbols—. Horizontal axis corresponds to the absolute value of the force modulus applied along $\mathbf{a}_1$, force is applied towards $-\mathbf{a}_1$. Simulation time 1.6 s. B) Mean lateral velocity for an ensemble of $10^3$ independent trajectories simulated in a landscape Eq. (2) with asymmetries $a_1/l_1 = 0.24$, $a_2/l_2 = 0.47$—blank symbols—, and $a_1/l_1 = 0.24$, $a_2/l_2 = 0.40$—black filled symbols—. Horizontal axis corresponds to the absolute value of the force modulus applied along $\mathbf{a}_2$, force is applied towards $-\mathbf{a}_2$. C) Rotational pitch and D) frequency of rotation for an ensemble of $10^3$ independent trajectories at zero load force in a landscape generated by Eq. (2) with asymmetry $a_1/l_1 = 0.24$, $a_2/l_2 = 0.47$ and varying dwell times in the range of $\tau^* = 36 - 72$ ms, in 4 ms steps.

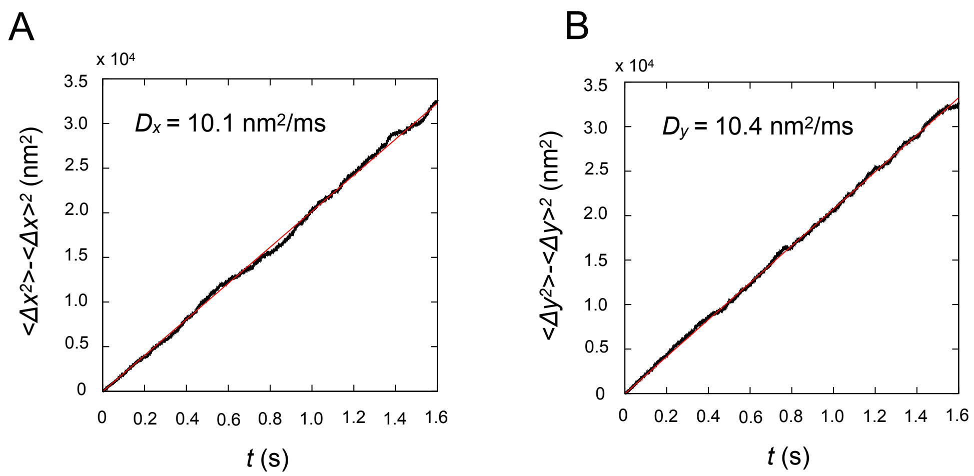


Supplementary Figure 5: Variance along the transversal (A) and longitudinal (B) axis obtained using an ensemble average with $10^3$ KIF1A trajectories and the landscape generated by Eq. (2) with $\mu_{11} = 1$, $\mu_{12} = 0.9$, $\mu_{13} = 0.65$, $\mu_{14} = 0.35$, $\mu_{21} = 1$, and $\mu_{22} = 0.4$.
